# Comprehensive Network Analysis and Experimental Verification of Dedu Safflower’s Influence on Ascites Hepatocarcinoma

**DOI:** 10.1101/2020.02.17.952085

**Authors:** Lu Wang, Sha Li, Lidao Bao

## Abstract

**Background:** Dedu Safflower Powder is a kind of classical prescription of Mongolian Medicine, and its main ingredients are the safflower and the Scabiosa tschiliensis. In the former Mongolian Medicine clinical practice, Dedu Safflower Powder, etc. had obvious effect on curing hepatocarcinoma so as to ease ascites. But the principle of Dedu Safflower Powder’s curing ascites hepatocarcinoma has been not clear.

**Aim:** H22 mouse hepatocarcinoma ascites models are used for researching the safflower and the Scabiosa tschiliensis’s function of jointly being against hepatocarcinoma ascites, and for verifying their possible combination principle (miR-23a-DAPK1-PVT1 signal axis).

**Method:** Through simulating an interaction network of the safflower and the Scabiosa tschiliensis with target spots, the function target was predicted. H22 mouse hepatocarcinoma ascites models were randomly divided into a blank contrast group, a model contrast group, a safflower group, a Scabiosa tschiliensis group and a compatibility group of the safflower/the Scabiosa tschiliensis. On the 10^th^ day, mice were killed to measure their weights and abdominal perimeters, also to collect ascites and blood for physical examinations. Kidney tissues were dissected at once and fixed in paraformaldehyde, for a real-time quantitative polymerase chain reaction (qRT-PCR) analysis.

**Result:** In the network, the DAPK1-PVT1 interaction showed the biggest edge betweenness, so it was predicted that DAPK1 and PVT1 were respectively presumed targets of the safflower and the Scabiosa tschiliensis. Compared with the model contrast group, the safflower group, the Scabiosa tschiliensis group and the safflower/Scabiosa tschiliensis group all had decreasing ascites volumes, body weights, abdominal perimeters. Compared with the model group, the safflower group and the safflower/Scabiosa tschiliensis group had rising DAPK1 expressions (p<0.05), while the safflower group had more obvious increasing DAPK1 (p<0.01); the Scabiosa tschiliensis group and the safflower/Scabiosa tschiliensis group had lowering PVT1 expressions (p<0.05), while the Scabiosa tschiliensis group had more obvious lowering PVT1 (p<0.01); the safflower group, the Scabiosa tschiliensis group and the safflower/Scabiosa tschiliensis group had lowering miR-23a expressions (p<0.05), while the safflower/Scabiosa tschiliensis group had more obvious lowering (p<0.01).

**Conclusion:** DAPK1 and PVT1 are respectively the presumed targets of the safflower and the Scabiosa tschiliensis. So Dedu Safflower Powder has good effects on ascites hepatocarcinoma, and that function may be partly related to regulation of miR-23a-DAPK1-PVT1 signal axis.

## Background

Hepatocellular Carcinoma (HCC) is one of the most common malignant tumors of the digestive system, and the ascites is one of complications of an advanced hepatocarcinoma, which refers to abnormal liquid accumulation in the abdominal cavity brought by tumor infiltration or secretion often worsening patients’ life quality^[1]^. At present, there are relatively much clinical treatment, but curative effects are not satisfactory, so malignant ascites treatment is still difficult. Traditional Chinese national medicine has natural advantages in controlling HCC patients’ condition development, improving symptoms and signs and raising living quality, etc.^[2]^.

Mongolian Medicine theory sets the concept of wholism as guidance, mainly led by simple materialism and spontaneous syndrome differentiation^[3]^. In Mongolian Medicine, there is a precedent of using the safflower and the Scabiosa tschiliensis to cure liver cancers so as to cure malignant ascites, because the safflower (HH) is feverfew, and its dry tubular flowers have effects of invigorating the circulation of blood, restoring the menstrual flow, removing blood stasis and relieving pains, having been used for hundreds of years in curing the ascites and hepatitis, etc.^[4]^. The Scabiosa tschiliensis (LPH) is a dipsacaceae plant, and its dry inflorescence is one of the common medicines for curing hepatic heat. The safflower and the Scabiosa tschiliensis have been used for curing hepatopathy so as to relieve ascites in the classical Mongolian Medicine prescriptions, which means compatibility of those two medicines may cause synergistic reaction^[5]^.

Network pharmacology, based on a interaction network of “diseases,genes, targets, medicines”, comprehensively and systematically observes medicine’s effects and interventions on diseases, so as to reveal the secret of medicines synergistic reaction on human bodies^[6]^. Consistent with the multi-ingredient, multi-channel and multi-target synergistic reaction principle of Mongolian Medicine and its prescriptions^[7]^, according to network pharmacology, the interrelationship of compatibility of the safflower and the Scabiosa tschiliensis on the objective network have been simulated, with explaining clearly relevance between the medicine target network topology and the medicine compatibility effect.

## Material and Method

### Structure Information of Chemical Composition of the Safflower and the Scabiosa tschiliensis

Traditional Chinese Medicine Chemistry Database (http://tcm.cmu.edu.tw /, updated on June 28, 2017), used for checking and collecting structure information of the safflower and the Scabiosa tschiliensis^[8]^.

### Prediction of HH and LPH Presumed Objective

HH and LPH presumed objectives were predicted by medicine similarity search tools at the target treatment database (TTD, http://xin.cz3.nus.edu.sg/group/cjttd/ttd.asp, version 4.3.02), and similar medicines to HH and LPH were selected by structure similarity comparisons.^[9]^.

### Network Construction and Analysis

HH and LPH presumed objectives, known liver cancer treatment targets and known treatment target for other human proteins were used for constructing medicine target PPI (protein-protein interaction) network. Through calculating such four topological characteristics (the bigger the value is, the more important the node is) as“degree” (the line amount for link nodes), “node betweenness” (proportion of route amount of passing that node to shortest route gross), “compactedness” (sum of distances of a node from all other nodes) and “K value” (the value for measuring node centrad), HH and LPH’s main presumed objectives were confirmed^[10]^.

### Molecular Abutment Simulation

DAPK1 and PVT1 were used respectively for molecular abutment simulation to direct bonding efficiency of main chemical ingredients of HH and LPH, with electronic high throughput screening (eHiTS) abutment module adopted^[11]^.

### U Medicine Preparation

HH (Batch Number: 160713) and LPH (Batch Number: 161106) were bought from Anhui Fengyuan Tongling Traditional Chinese Medicine Company Limited. HH was boiled out at 1.34 g/ kg stoste, and 20.1g HH were soaked in 10 times water for 1h, and then that was boiled for 1.5h before using a electrotherm to heat, and after stewing and filtering, residuals were boiled for 1h by water of the same amount. After again stewing and filtering, filter liquor was merged, and then evaporated and condensed it to a specified volume (300ml) at 60°C, and then it was kept at 4°C. LPH were pulverized by a grinder, and then filtered by an 80-eye screen. Those powder was left for the next step. LPH powder and HH apozem (stoste) were mixed at 1:1 and heated before the intragastric administration of giving medicine.

### Cell Cultivation and Animal Model

Mouse H22 hepatocarcinoma ascites cell lines were bought from Clinical Experiment Center of Inner Mongolia Medical University. H22 cells were cultivated in the rpmi - 1640 culture medium, supplemented with 2mm l-glutamine, 100iu / mL penicillin, 100ug/ml streptomycin and FCS below 10%, at 37 °C with wet atmosphere containing 5%CO_2_, with one generation cultivated every 2-3 days. 18-22g male BALB / C mice (6-week-old) were acquired from Laboratory Animal Centre of Inner Mongolia Medical University, and they were left to adapt to new environments one week before the experiment, with all mice raised in a laminar flow cabinet condition of no causative agents. H22 liver cancer ascites tumor models were prepared. Needles were inserted into the left lower quadrant, so the abdominal cavity was injected with H22 cells (except the blank contrast group). Every mouse was injected with 1 × 10^7^ H22 cells, and that operation was not related to mortality or incidence.

### Grouping and Treatment

In this research, all H22 hepatocarcinoma ascites models were divided into five groups (10 mice for every group): (1) blank contrast group, (2) model contrast group, (3) safflower group (0.93g/kg), (4) Scabiosa tschiliensis group (1.02g/kg), (5) safflower/Scabiosa tschiliensis combination group (HH/LPH ratio is 1/1, namely 0.46g/kg safflower and 0.51g/kg Scabiosa tschiliensis).

H22 hepatocarcinoma ascites mice in HH group, LPH group and HH/LPH combination group were respectively given intragastric administration of the above HH and LPH of proper dosages. In the blank contrast group and the model contrast group, mice were given normal saline of the same volume. Mice for experiment were raised in a standard lab condition, and used for different experiments.

### Physical Check

On the 10^th^ day, living mice were killed by cervical dislocation, to measure mice’s weights and abdominal perimeters. The ascites was collected from the open napes of cervical dislocation, and was measured by a injector.

### Fluorescence Timed Quantitative Polymerase Chain Reaction (qRT-PCR)

Killed mice’s kidney tissues were dissected and fixed in paraformaldehyde, to be done fluorescence timed quantitative polymerase chain reaction (qRT-PCR), with the experiment conducted according to conventions^[12]^. GAPDH was set as the target gene (Forward: 5’-TGAGACATGAAGGTTTCGAAG-3’Reverse: 5’-TCCTTATTTGGGGCTTGGCCAT-3’). DAPK1 primer sequence is Forward:5’-GCCTTCTGGCTCATTGCGATGATT G −3’; Reverse: 5’-CAGCAATCGGTGTGGTTGTGTTGT −3’. PVT1 primer sequence is Forward: 5’-TGCGGAGGATGACACTGAGTGC −3’; Reverse: 5’-TGTGCCTGCTACCAGAATTGCTC −3’. miR-23a primer sequence is Forward: 5’-AGAGCGATCCTTACGGTTCTGTG −3’. All experiments wer in triplicate.

### Statistical Analysis

SPSS software of the version 16 (SPSS Company, Chicago, IL, America) and SAS 9.1 (SAS Institute, Kari, NC) were used for a statistical analysis. The continuous variable was expressed as mean±standard deviation, and many group data were compared, with the way analysis of variance and LSD test used. Difference is of statistical significance, P < 0.05.

## Result

### Mongolian Medicine HH and LPH Medicine Target

According to Traditional Chinese Medicine Chemical Database, there were 89 HH chemical compounds and 10 LPH chemical compounds. HH and LPH presumed objectives are respectively 14 and 96. HH and LPH target PPI (protein interaction) and other known human protein treatment targets were used for building up a medicine target mesh which has 3955 nodes and 7735 sides, and if a node’s degree is more than two times mid-values of all nodes, that can be regarded as a center. Thus, 636 nodes were confirmed as the center. We have constructed an interactive network of those centers,consisting of 463 nodes and 1976 sides, just as Picture 1A. DAPK1 and PVT1 are respectively main presumed objectives of HH and LPH, having interreaction with miR-23a in functions, so we think that HH/LPH combination influence on hepatocarcinoma ascites may be related to miR-23a-DAPK1-PVT 1 signal axis, just as the Picture 1B.

### Molecular Abutment Simulation Result and Physical Examination

eHiTS software was used for molecular abutment simulation, and the higher absolute values of abutment grades,the stronger direct binding efficiency of the ligand and the receptor is. According to the marking system of eHiTS algorithm, in 89 chemical ingredients of HH, 60 (66.67%) have medium-strong joint efficiency with DAPK1, while in 30 chemical ingredients of LPH, 20 (66.67%) have medium-strong joint efficiency with PVT1.

Compared with the model contrast group, the HH group, the LPH group and the HH/LPH group all have reduced ascites volumes (Picture 2A), weights (Picture 2B) and abdominal perimeters (Picture 2C). Difference of the HH/LPH group and the model contrast group is more obvious than that of the HH group, the LPH group and the model contrast group (Picture 2).

### DAPK1 Expression Functions in Ascites, Serum and Kidney Tissue

We have confirmed that DAPK1 is the main presumed objective of HH. In ascites, serum and kidney tissues of the HH group, DAPK1 protein and mRNA expression levels were obviously higher than the model control group (P<0.01). The model control group had difference with the HH/LPH compatibility group (P<0.05), but no obvious difference with LPH group (Picture 3).

### PVT1 Expression Function in Ascites, Serum and Kidney Tissue

We have confirmed that PVT1 is the main presumed objective of LPH. PVT1 protein and mRNA expression levels in ascites, serum and kidney tissues in the LPH group were obviously lower than the model control group (P<0.01). The model control group had difference with the HH/LPH compatibility group (P<0.05), but no difference with the HH group (Picture 3).

### miR-23a Expression Function in Ascites, Serum and Kidney Tissue

miR-23a is the known gene causing liver cancers. miR-23a levels of HH group and LPH group in ascites, serum and kidney tissues had difference with the model contrast group (P<0.05), while expression levels of ascites, serum and kidney tissues in the HH/LPH compatibility group were obviously lower than model contrast group (P<0.01) (Picture 5).

## Discussion

Based on the four topological characteristics of the interaction network of the network Pharmacology Center, we use Markoff clustering algorithm to divide the main presumed objective interaction network into 3 functional modulest^[13]^. According to the enrichment analysis of the GO annotation system and the KEGG path, side betweenness of every interaction in the main presumed objective network were calculated. Interaction between DAPK1 and PVT1 shows the biggest boundary value, indicating its importance in connecting different modules in the network^[14]^.

DAPK1 is a serine / threonine protein kinase regulated by 160 k U calmodulin (Ca M), a positive regulatory factor of apoptosis closely related to tumor occurrence, development and metastasis^[15]^. DAPK1, as a anti-oncogene joining the cell apoptosis, plays an important role in many routes guiding cell apoptosis, such as the cell apoptosis guided by TNF α, the cell apoptosis guided by related suicidal factors, the cell apoptosis guided by interferon y and he cell apoptosis guided by TGF β^[16]^ HH has obvious up-regulation function on DAPK1, and because HH content of the HH/LPH compatibility group was lower than the HH group, DAPK1 levels difference in ascites, serum and kidney tissues in the model control group with the HH group were higher than difference between the model contrast group and HH/LPH compatibility group^[17]^.

The human PVT1 gene, also known as plasma-cytoma variant trans-location gene 1, has length more than 300kb, also the PVT1 gene is located in the chromosome 8q24 area and in the positive-sense strand of the chromosome, besides chromosome 8q24 area is the highest target of DNA copy number amplification, whose abnormal amplification often implies a high tumor onset risk^[18]^. According to research in recent years, PVT1, as a tumor specific gene, plays an important role in such many tumors as prostate cancer,lung cancer, liver cancer, gastric cancer and colorectal cancer, joining such biology functions as tumor cells’ proliferation, apoptosis, invasion and metastasis, stem cell differentiation and medicine resistance. Its expression in HCC increases obviously^[19]^. LPH has some certain regulation function on PVT1. However, PVT1 levels difference in ascites, serum and kidney tissues of the model control group with the LPH group was higher than model contrast group difference with the HH/LPH compatibility group. Because the LPH dosage in the HH/LPH compatibility group is so low, intensity of regulating down its presumed objectives is not hight^[20]^.

miR-23a is a member of miRNA. Resent research has shown that miR-23a widely regulates and controls every physiological and pathological process of the body. miR-23a plays an important role in cell differentiation, joining such processes as skeletal muscle forming, hematopoietic cell differentiation, oligodendrocyte differentiation, etc.; has a function of causing cancer in liver tumors. miR-23a and its target genes constitute a complex regulation and control network, which plays an important role during cell proliferation, differentiation, damage, apoptosis and tumor formation as well as production of resistance to medicines^[21–22]^. HH and LPH all have functions of regulating down miR-23a. However, treatment effects of HH and LPH separate application were not obvious than those of joint application of those two, indicating that the combination of two medicines can resist ascites hepatocarcinoma. Moreover, it has been found in our network analysis interaction between DAPK1 and PVT1, indicating that signal axis of miR-23a-DAPK1-PVT1 can weaken the hepatocarcinoma ascites^[23]^.

## Conclusion

This research combined the medicine objective prediction, network analysis with experimental verification, supporting convincing evidence of that HH and LPH compatibility can resist ascites hepatocarcinoma, which may be partly related to their laws of function on the miR-23a-DAPK1-PVT1 signal axis.

## Acknowledgement

This research is funded by the following projects: National Natural Science Foundation of China (No. 81760748 and 81550047); the Major Science Foundation of Affiliated Hospital of Inner Mongolia Medical University, Hohhot, Inner Mongolia.

---

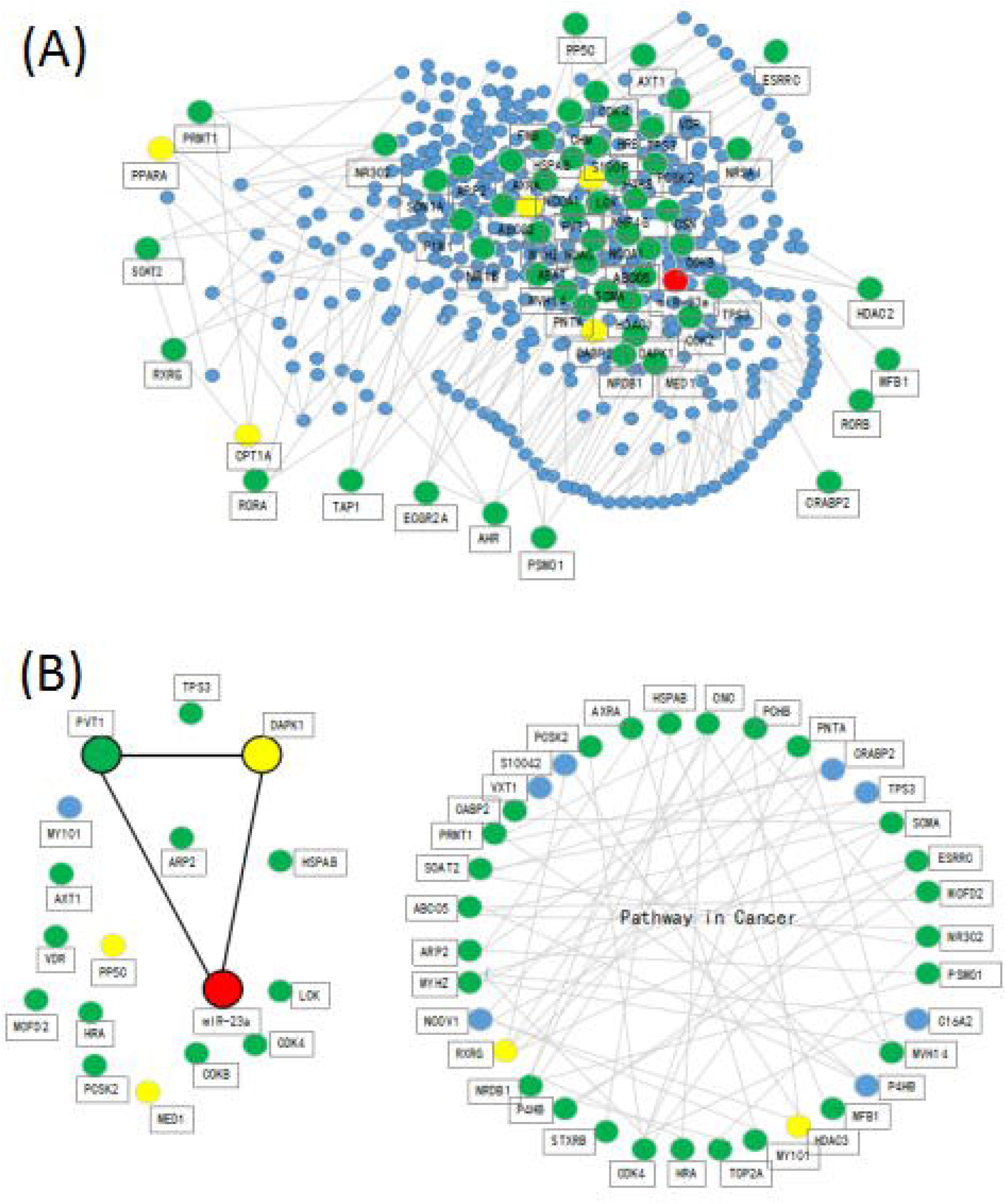

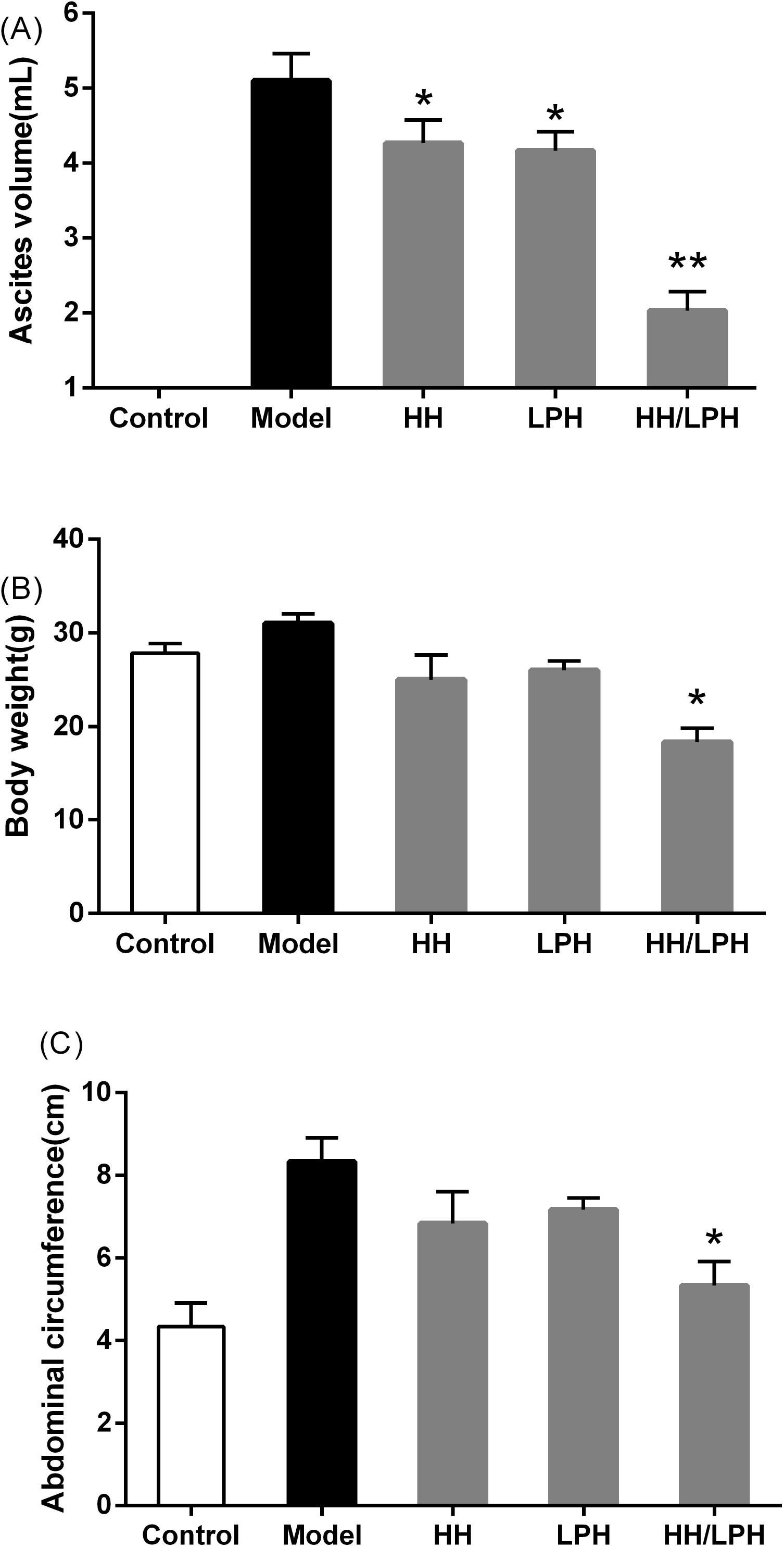

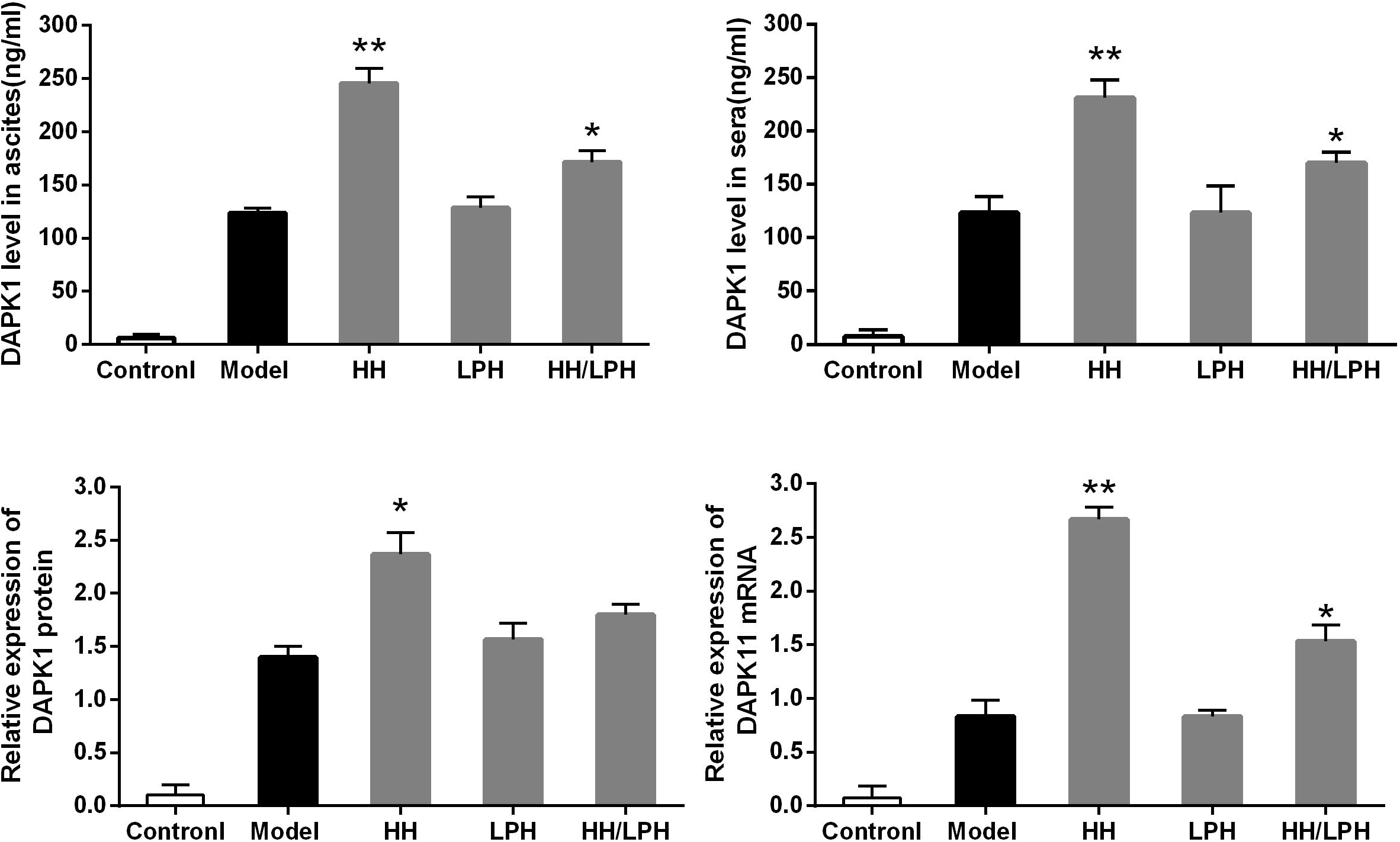

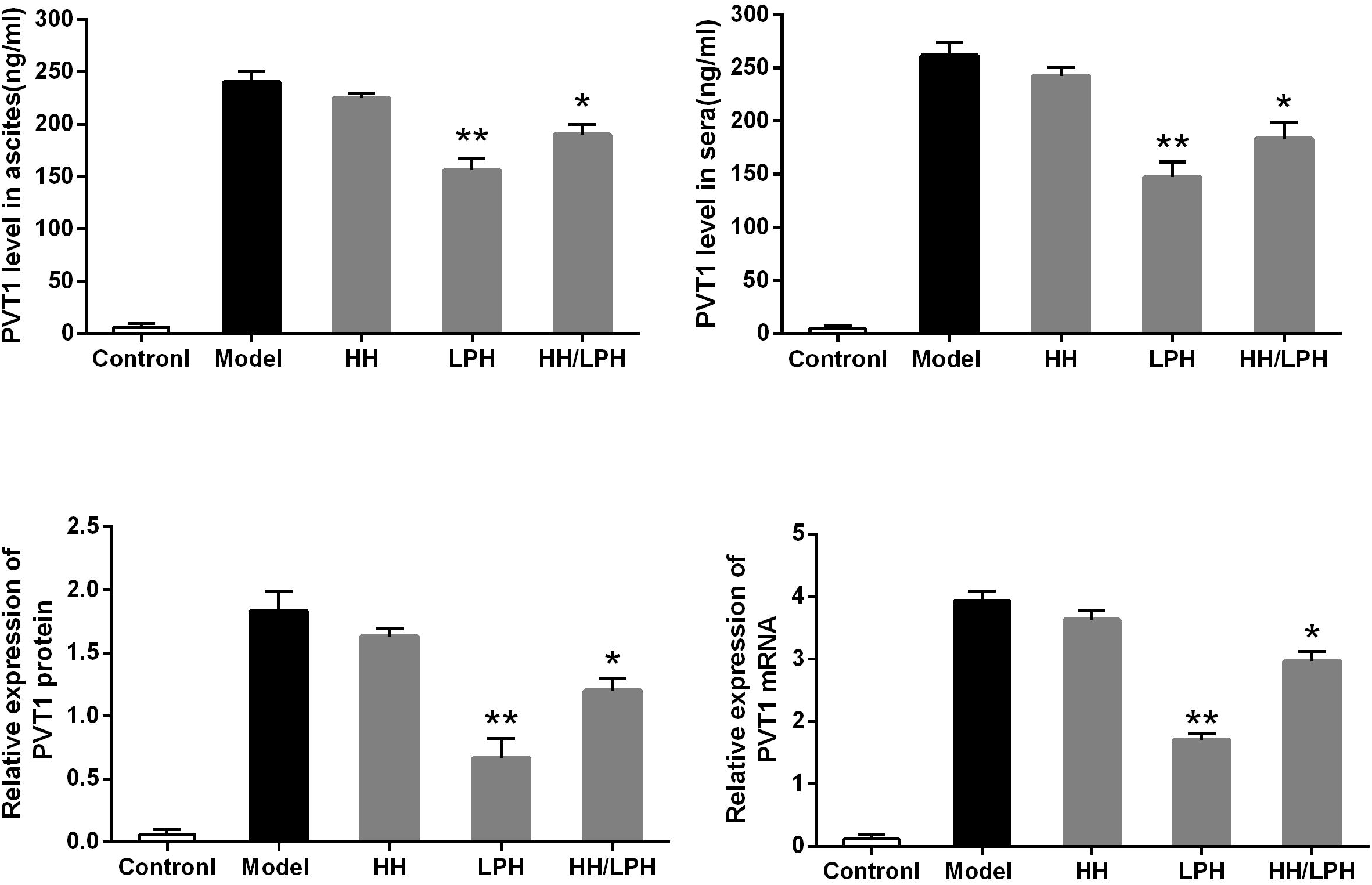

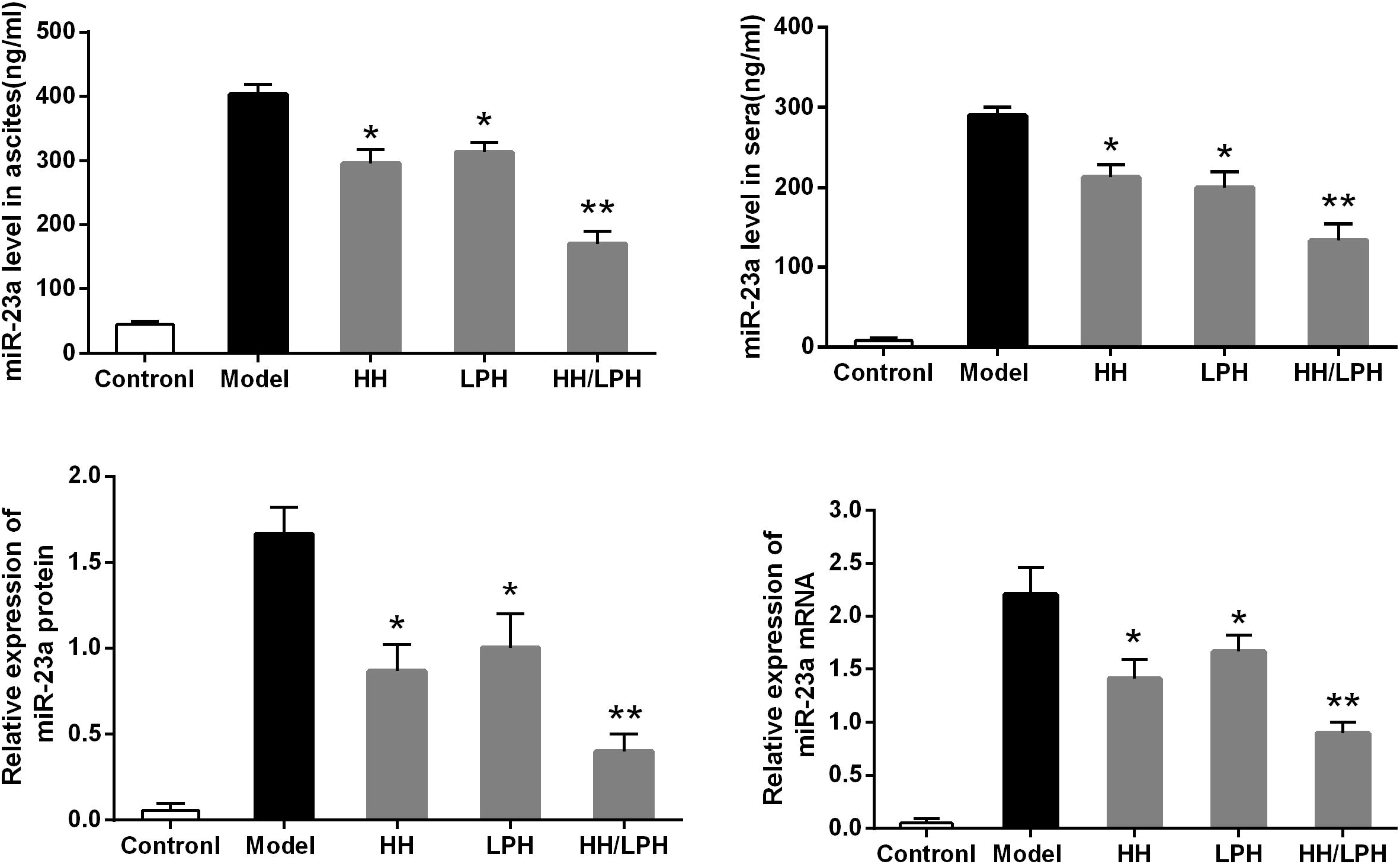

